# Middle-level IM-MS and CIU experiments for improved therapeutic immunoglobulin isotype fingerprinting

**DOI:** 10.1101/2020.01.20.911750

**Authors:** Thomas Botzanowski, Oscar Hernandez-Alba, Martine Malissard, Elsa Wagner-Rousset, Evolène Deslignière, Olivier Colas, Jean-François Haeuw, Alain Beck, Sarah Cianférani

## Abstract

Currently approved therapeutic monoclonal antibodies (mAbs) are based on immunoglobulin G (IgG) 1, 2 or 4 isotypes, which differ in their specific inter-chains disulfide bridge connectivities. Different analytical techniques have been reported for mAb isotyping, among which native ion mobility mass spectrometry (IM-MS) and collision induced unfolding (CIU) experiments. However, mAb isotyping by these approaches is based on detection of subtle differences and thus remains challenging at the intact-level. We report here on middle-level (after IdeS digestion) IM-MS and CIU approaches to afford better differentiation of mAb isotypes. Our method provides simultaneously CIU patterns of F(ab’)_2_ and Fc domains within a single run. Middle-level CIU patterns of F(ab’)2 domains enable more reliable classification of mAb isotypes compared to intact level CIU, while CIU fingerprints of Fc domains are overall less informative for mAb isotyping. F(ab’)_2_ regions can thus be considered as diagnostic domains providing specific CIU signatures for mAb isotyping. Benefits of middle-level IM-MS and CIU approaches are further illustrated on the hybrid IgG2/IgG4 eculizumab. While classical analytical techniques led to controversial results, middle-level CIU uniquely allowed to face the challenge of eculizumab « hybridicity », highlighting that its F(ab’)_2_ and Fc CIU patterns corresponds to an IgG2 and an IgG4, respectively. Altogether, the middle-level CIU approach is more clear-cut, accurate and straightforward for canonical but also more complex, engineered next generation mAb formats isotyping. Middle-level CIU thus constitutes a real breakthrough for therapeutic protein analysis, paving the way for its implementation in R&D laboratories.

## INTRODUCTION

During the last 20 years, monoclonal antibody (mAb) development and engineering have significantly evolved due to their therapeutic efficiency against many diseases such as cancer, and autoimmune diseases^1^. More than 80 antibody-based products are currently approved by regulatory agencies (FDA and EMA), while ~600 others are in clinical studies, including more than 60 in phase III clinical trials^2^.

While human IgG3 subclass is usually not considered for therapeutic mAbs engineering and production due to its limited potential associated to its shorter half-life^1^, human IgG1, IgG2, and IgG4 mAb isotypes represent the main classes of mAb-based therapeutics. One of the main structural differences between the three therapeutic isotypes (IgG1, IgG2, and IgG4) is the number (four for IgG1 and IgG4 and six for IgG2) and the connectivities of inter-chain disulfide bridges^3^ (Figure 1). The heavy and light chains of all isotypes are linked by one disulfide bond, while the two heavy-chains can be linked either by two (for IgG1 and IgG4) or four (for IgG2) disulfide bonds located in the hinge region of the antibody^3^. In addition, mAbs global structure is also maintained with 12 intra-chain disulfide bridges that connect two cysteines that belong to the same domain. The inter-chain disulfide bridges network, which is characteristic of each individual isotype, has an impact on different mAb properties (structure, stability, surface hydrophobicity, isoelectric point, etc.)^4^ and modulate their higher-order structure^5–8^. Thereby, mAbs from different isotype classes will differ in their secondary immune functions^9–10^. In terms of mAb developability, there is a continuous interest for improvement of new analytical techniques to characterize the impact of the different inter-chain disulfide patterns on therapeutic mAb structures and structure-function relationships.

**Figure 1:**
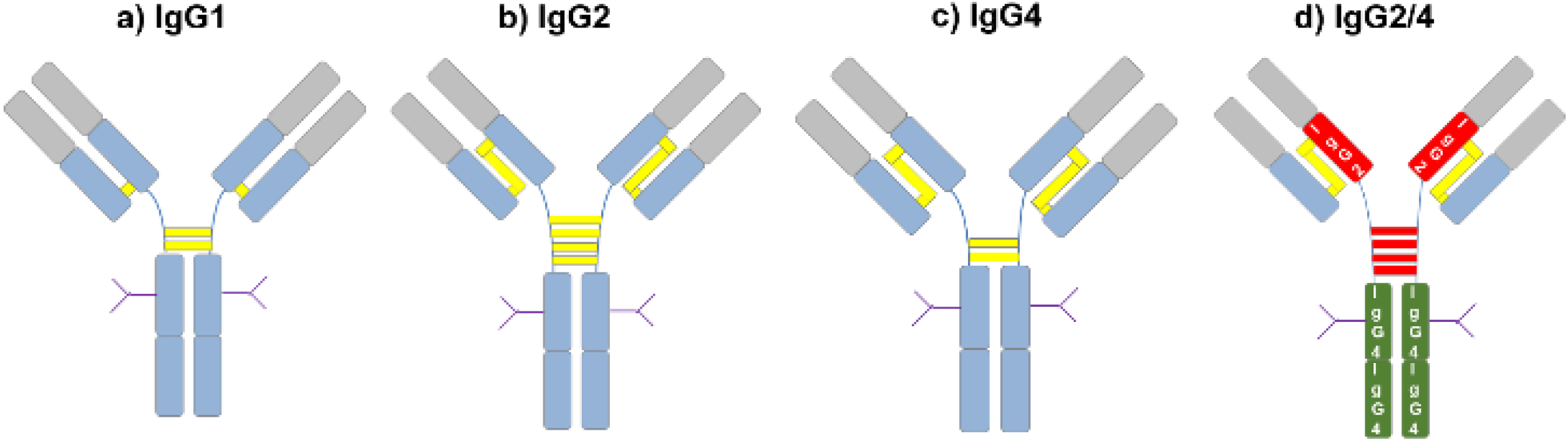
Schematic representation of inter-chain disulfide bridges (yellow bars) characteristic of the IgG1 (a), IgG2 (b), IgG4 (c) and hybrid IgG2/4 (d) mAb isotypes.

Ion mobility coupled to mass spectrometry (IM-MS), and its collision induced unfolding (CIU) variant, have been used to characterize the structure and dynamics of proteins^11–14^ and protein complexes^15–16^ in the gas-phase. During the last 5 years, CIU has been increasingly used in structural biology to characterize a wide range of biological systems and has entered the analytical portfolio of international regulatory agencies^17^. Although the CIU approach still remains as a laborious and time-consuming process, significant efforts have been made to improve data acquisition^18^ and interpretation^19–20^. CIU experiments allowed to circumvent in some cases the limitations associated to IM resolution to separate and differentiate mAbs with very similar global structure (less than 5 % of CCS difference). Thereby, CIU afforded structural insights that led to the differentiation of human non-therapeutic mAb isotypes^20–21^, ADCs’ characterization^22–23^, and stabilized vs wild-type therapeutic IgG4 mAbs among others^24^. Ruotolo and coworkers reported that the four human mAb isotypes give rise to different unfolding patterns upon collisions with the background gas, showing the influence of the inter-chain disulfide bridges on mAb gas-phase stability^21^. However, in some cases, the categorization/characterization of mAb isotypes remains challenging due to the very subtle differences observed in their intact CIU fingerprints.

In the present work, we aimed at improving IM-MS and CIU workflows to better differentiate the isotypes of therapeutic mAbs (IgG1, IgG2, and IgG4), including engineered hybrid mAb formats. For this purpose, we developed middle-level IMMS and CIU approaches where the mAb scaffold is IdeS-digested^25^ prior to IM-MS or CIU analysis. In this case, a thorough characterization of therapeutic mAbs scaffold is performed based on the individual analysis of the F(ab’)_2_, and Fc subdomains. The global structure along with the gas-phase dynamics associated with each subunit highlighted the structural similarities/differences induced by the inter-chain connectivities of each therapeutic isotype class, and provided clear-cut evidences to improve mAb isotypes differentiation. Finally, the combination of middle-level IM-MS and CIU also allowed to unravel the isotype of a hybrid engineered mAb, pinpointing the suitability of IM-MS and CIU at middle level to clearly characterize and differentiate the isotype of canonical and hybrid therapeutic mAbs of next generation therapeutics.

## MATERIALS AND METHODS

### Sample preparation

Eculizumab (Soliris, Alexion Pharmaceuticals Inc.), panitumumab (Vectibix, Amgen) natalizumab (Tysabri, Biogen), and adalimumab (Humira, Abbvie) were sourced from their respective manufacturers as EMA-approved drug products. Each individual mAb was N-deglycosylated during 30 min at 37 °C with IgGZERO (Genovis). In the case of middle-level analysis, the deglycosylated mAbs were degraded with IdeS enzyme (immunoglobulin-degrading enzyme of *Streptococcus pyogenes*, FabRICATOR, Genovis). A 1 µg/unit ratio was used to achieve an efficient digestion and subsequently, the mixture was incubated during 60 min at 37°C. After deglycosylation and/or IdeS digestion, therapeutic mAbs were then desalted against 100 mM ammonium acetate at pH 7.0 prior to native MS analysis, using about six to eight cycles of centrifugal concentrator (Vivaspin, 30 kDa cutoff, Sartorious, Göttingen, Germany). The concentration of each individual solution after desalting process was measured by UV absorbance using a nanodrop spectrophotometer (Thermo Fisher Scientific, France). Prior to native MS analysis, each sample was diluted in 100 mM ammonium acetate at pH 7.0 to a final concentration of 5 µM.

### Native MS analysis

Native mass spectra were acquired on an Orbitrap Exactive Plus EMR (Thermo Fisher Scientific, Bremen, Germany) coupled to an automated chip-based nanoelectrospray device (Triversa Nanomate, Advion, Ithaca, USA) operating in the positive ion mode. The capillary voltage and the pressure of the nebulizer gas were set at 1.7-1.9 kV and 0.15-0.20 psi, respectively. The source parameters were tuned in order to obtain best mass accuracy for native MS experiments as followed: briefly, the in-source voltage was set to 150 eV, the HCD cell voltage was fixed to 50 eV and the pressure of the backing region was fixed to 2 mbar. Native MS data interpretations were performed using Xcalibur^TM^ software v4.0 (Thermo Fisher Scientific, Bremen, Germany).

### Native IM-MS and CIU experiments

Ion mobility and CIU experiments were performed on a hybrid Q-IM-TOF mass spectrometer (Synapt G2, Waters, Manchester, UK). The cone voltage was fixed at 80 V to improve the ion transmission and avoid in source ion activation. The backing pressure of the Z-spray source was set to 6 mbar and the argon flow rate was 5 mL/min. Ions were cooled and separated in the IM cell with a Helium flow rate of 120 mL/min and a N_2_ flow rate of 60 mL/min. Ion mobility parameters were tuned to improve ion separation and prevent ion heating as described in *Hernandez et al*^24^. Briefly, the wave velocity and height were fixed to 800 m/s and 40 V, respectively. IM drift times of each mAb were converted in collision cross sections using three charge states of concanavalin A, pyruvate kinase and alcohol deshydrogenase as external calibrants. MassLynx software (Waters, Manchester, U.K.) was used to generate arrival time distributions. ^TW^CCS_N2_ values were calculated as an average of three replicates for mAb and calibrants under strictly identical experimental conditions.

Collision induced unfolding experiments were performed by increasing the collision voltage of the trap by 5 V steps from 0 to 200 V prior to IM separation. Individual IM data were gathered to generate CIU fingerprint using the CIUSuite2 software and in particular the CIUSuite2_BasicAnalysis and the CIUSuite2_StabilityAnalysis modules in order to obtain average and differential plots, and then to determine CIU50 values to assess the stability of each transition directly from the CIU data. Each plot corresponds to the average of the three analysis replicates with a root mean square deviation lower than 10 % showing a good reproducibility of the experiment. ATD intensities were normalized to a maximum value of 1 and classical smoothing parameters were used (Savitzky-Golay algorithm with a window length of 3 and a polynomial order of 2) as used in the previous version of the software. For isotype classification, we used three CIU replicates of adalimumab (IgG1 reference), panitumumab (IgG2 reference), and natalizumab (IgG4 reference). A statistical method based on the variance analysis (ANOVA) F-test was used to analyze the CIU fingerprints of eculizumab and assess the significance of the energy values for the isotype differentiation.

### RPLC analysis

Separation of the different IgG1, IgG2, IgG2/4 and IgG4 isotypes were performed in a Zorbax RRHD column (2.1 mm x 50 mm, 1.8 µm, 300 Å) from Agilent Technologies (Wilmington, DE, USA). The column was loaded with 1 µl of the intact mAbs solution at 5 mg/ml final concentration (5 µg). Mobile phase A was composed of 0.1% TFA, 2% isopropyl alcohol (IPA) in water, and mobile phase B was 0.1% TFA, 25% acetonitrile, in IPA. Samples were eluted with a constant flow rate of 250 µL/min and using a chromatographic gradient from 10 to 25% B over 9 minutes, followed by a shallow gradient up to 27.8% B over 7 min. Then, the gradient increased up to 29.8% B over 1 minute, followed by 29.8 – 50% B over 2 minutes.

### nrCE-SDS analysis

IgGs were analyzed in non-reduced condition using a Maurice^TM^ system (Protein Simple) equipped with the Compass^TM^ software. Chemicals were provided from the Maurice^TM^ CE-SDS application kit from the provider. Samples were diluted in 1x sample buffer to a final concentration of 1 mg/mL, from which 50-µL-aliquot samples were made. Then 2 µl of internal standard was added to each sample. 2.5 µl of a 250-mM stock solution of the alkylating agent iodoacetamide was added to each 50-µL sample to block disulfide scrambling or exchange. They were denatured at 70 °C for 10 minutes, cooled on ice for 5 minutes and mixed by vortex. Each sample was then transferred to a 96-well plate and spun down in a centrifuge for 10 minutes at 1000 x g. All samples were electrokinetically injected into the cartridge capillary by applying 4600 V for 20 seconds before separation by electrophoresis at 5750 V during 35 min. Electropherograms were analyzed with the Empower^TM^ data software.

### Differential scanning calorimetry (DSC)

DSC experiments were performed on a MicroCal VP-Capillary DSC instrument (Malvern Instruments). Samples were buffer exchanged into PBS Dulbecco pH 7.4 buffer or 25 mM His/His-HCl, 150 mM NaCl, pH 6.5 and diluted to 1 mg/mL in according buffer. 400 μL of the protein solution as well as 400 µL of the according buffer were dispensed in 96 well plates, loaded to the capillary sample cell while the reference cell contained the corresponding buffer. The chamber was pressurized to 3 atm and the temperature ramped from 40°C to 100°C at 1°C/min heating rate. The recorded DSC thermograms were baseline subtracted and subjected to a multi-component Gaussian fitting in the MicroCal VP-Capillary DSC software 2.0 (Malvern Instruments).

The temperatures for three major transitions were extracted from the fitted Gaussian models, relating to the unfolding of CH2, Fab, and CH3 domains. For each sample, 3 independent experiments were carried out allowing us to use a value of 1°C as the cutoff limit for evaluating the significance of the differences observed in melt temperatures.

## RESULTS AND DISCUSSION

### Intact level IM-MS and CIU experiments for mAb isotype classification

We first analyzed three therapeutic mAbs - adalimumab (IgG1), panitumumab (IgG2), and natalizumab (IgG4) – at the intact level by native MS (Figure S1). Overall, the charge state distributions (CSD) observed on the native mass spectra are centered either on the 24+ or 23+ charge states. The same therapeutic mAbs were next analyzed by native IM-MS at the intact level to provide more insights into their global conformation (Figure S2). For all charge states, IM-MS provide very similar ^TW^CCS_N2_ values within the error of the CCS measurement, avoiding classification of isotypes on the sole basis of CCS measurements. The co-elution of all reference therapeutic mAbs upon IM separation leads to the conclusion that current TWIMS resolution cannot afford an efficient differentiation of the three isotypes as previously reported on non-therapeutic mAbs^21, 24^ (Figure S2a). The high similarity in terms of primary sequence between these mAbs leads to the analysis of quasi-isobaric (< 2 % mass difference) and quasi-iso-cross sectional (< 3 % CCS difference) (Figure 2b, c) proteins for which classical native IM-MS instrumentation can only provide limited information.

**Figure 2:**
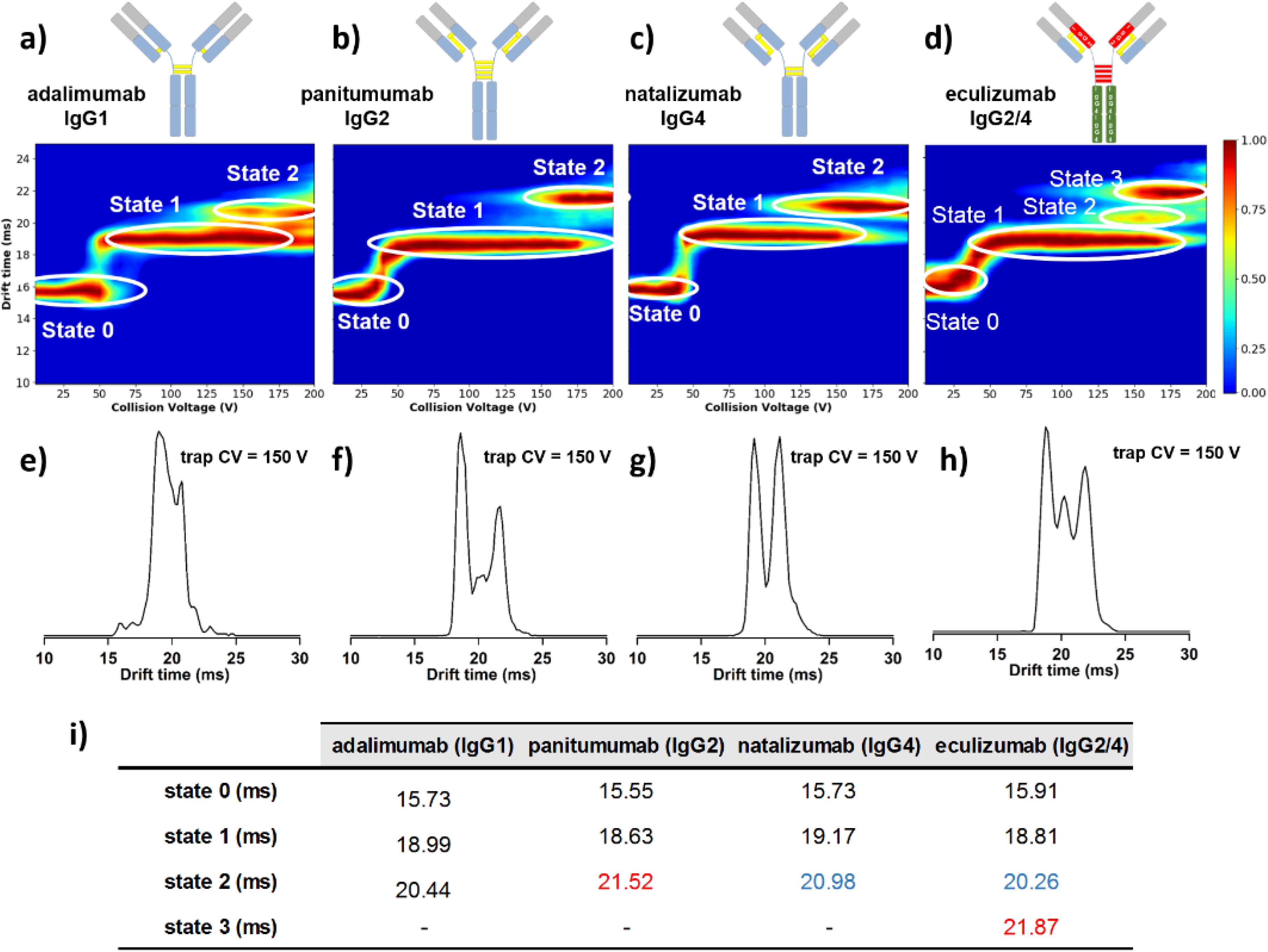
Intact level CIU experiments. CIU experiments of the 22+ charge state of adalimumab (IgG1) (a), panitumumab (IgG2) (b), natalizumab (IgG4) (c), and hybrid IgG2/4 eculizumab (d). CIU fingerprints are depicted in the upper panels. ATDs extracted at 150 V corresponding to the three therapeutic mAbs are depicted in the lower panels (e, f, g, and h). Table summarizing the IM drift times of the observed unfolding states (i).

As CIU was reported to be sensitive to the number and pattern of inter-chain disulfide bridges contained within the structure of mAbs from human serum and thus leading to CIU fingerprints characteristic of each mAb isotype^20–21^, we next performed and compared CIU experiments on the three therapeutic mAbs at the intact level (Figure 2). Overall the CIU patterns of the three mAb isotypes look very similar with two unfolding transitions present on the three CIU fingerprints. While the three canonical mAb isotypes (adalimumab, panitumumab, and natalizumab) exhibit the same IM migration times at the ground state, some subtle differences can be observed upon ion activation (Figure 2a, b, and c). The most and unique diagnostic CIU region is comprised between 100-200 V range where ATDs from the three isotypes exhibit different distributions (Figure 2e), allowing the discrimination of the therapeutic mAb isotypes at the intact level, as previously reported^20–21, 24^. Although mAb isotypes can be differentiated when these structures populate excited unfolding states upon activation with the background gas, CIU fingerprints at the intact level only provides very limited and subtle differences, hindering a clear-cut classification of therapeutic mAb isotypes.

### Middle-level IM-MS and CIU analysis for clear-cut mAb isotype classification

In order to circumvent intact level IM-MS and CIU limitations, we next performed native IM-MS and CIU experiments (Figure 3, 4, and 5) at the middle level to further characterize the global conformation and the gas-phase stability of the F(ab’)_2_ and Fc domains of IdeS-digested adalimumab, panitumumab, and natalizumab (see Material and Method section). For the 20+ and 21+ charge states, the measured ^TW^CCS_N2_ of the (Fab’)_2_ domains allow clear differentiation of IgG2 isotype from IgG1/IgG4 but unables distinguishing IgG1 from IgG4 which show similar ^TW^CCS_N2_, probably due to the same number and very close intra-chain disulfide bridge connectivities in the hinge region (Figure 3). For the 12+ charge state of the Fc region, the measured ^TW^CCS_N2_ are 33.1 ± 0.1 nm², 33.2 ± 0.1 nm² and 33.1 ± 0.1 nm² for IgG1, IgG2 and IgG4 references respectively (Figure S3), showing that the categorization/characterization of mAb isotypes cannot be performed based on the IM data of the Fc domains.

**Figure 3:**
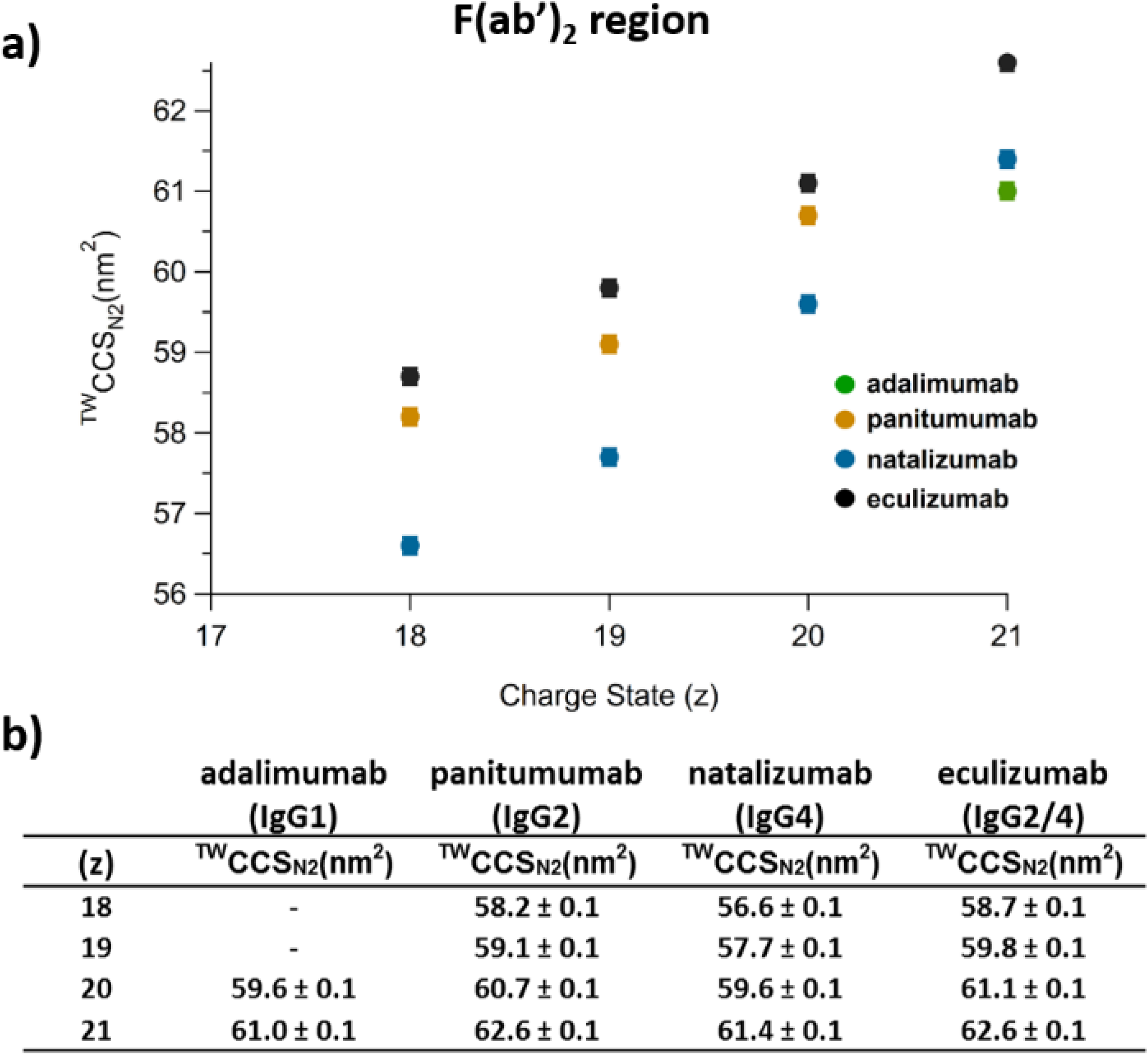
Middle-level IM-MS analysis of F(ab’)_2_ domains. Evolution of the F(ab’)_2_ ^TW^CCS_N2_ as a function of the charge state (a). Table summarizing the measured ^TW^CCS_N2_ of the F(ab’)_2_ domains.

**Figure 4:**
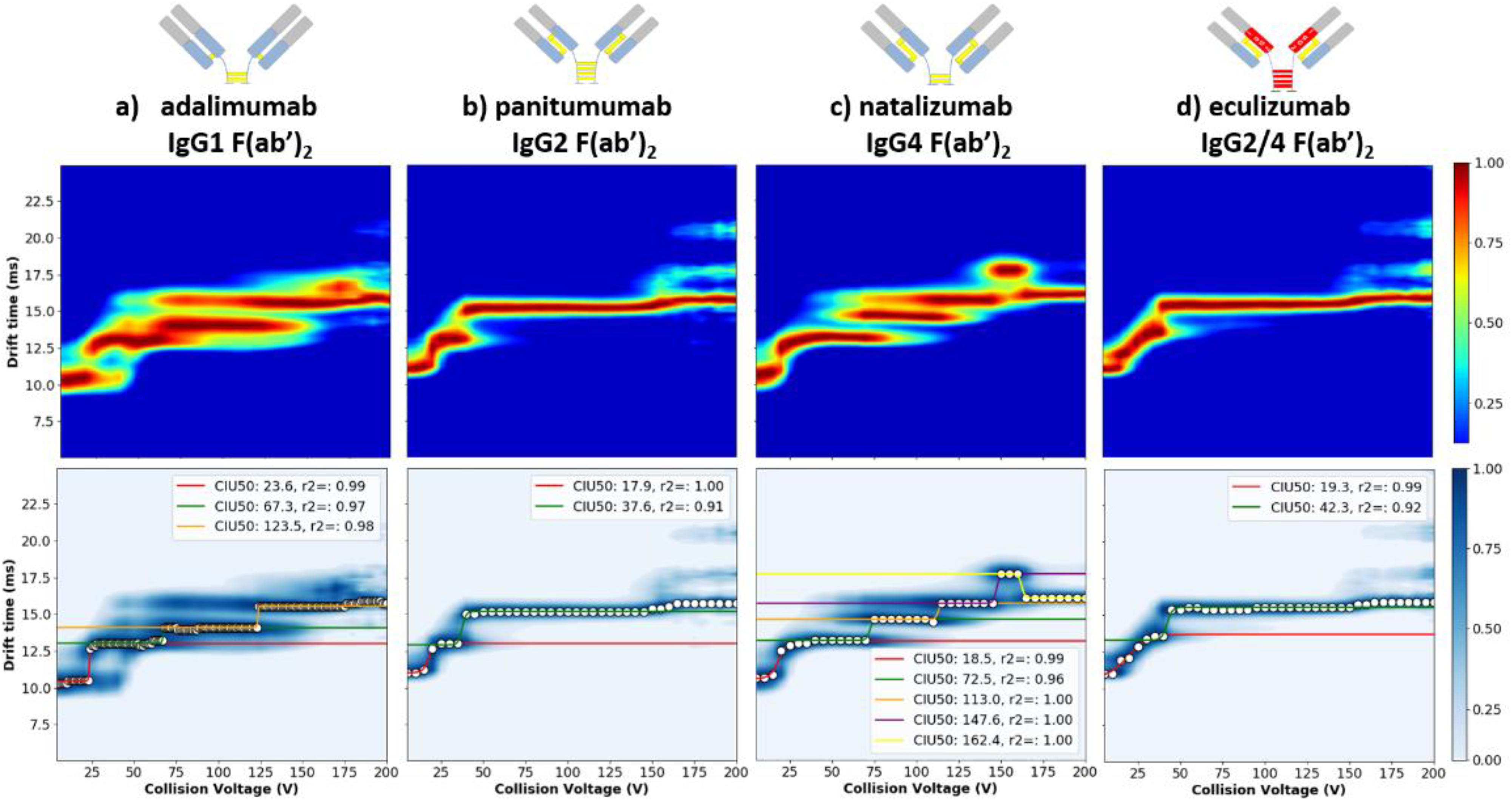
Middle-level CIU experiments on F(ab’)_2_ domains. CIU fingerprints (top panel) and stability analysis “CIU50” of 21+ charge state of F(ab’)_2_ domain of IgG1(a), IgG2 (b), IgG4 (c), and IgG2/4 (d) from 0 to 200 V trap collision voltage. Gaussian fitting and collision voltages associated with the unfolding transitions are depicted in the lower panels.

**Figure 5:**
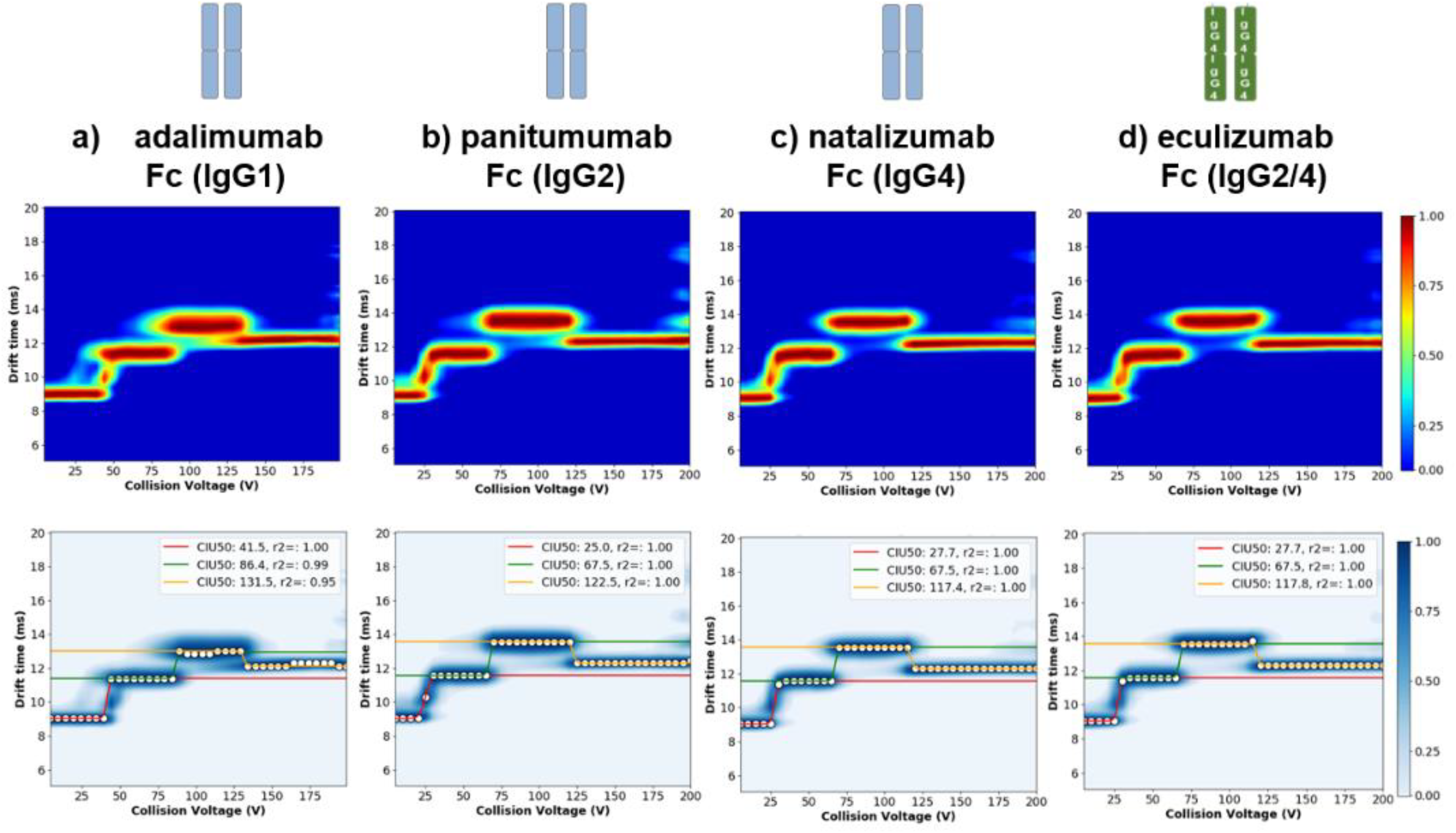
Middle-level CIU experiments on Fc domains. CIU fingerprint of 12+ charge state of Fc domains corresponding to adalimumab (IgG1) (a), panitumumab (IgG2) (b), natalizumab (IgG4) (c), and eculizumab (IgG2/4) (d). The CIU fingerprints and the corresponding Gaussian fitting are depicted in upper and lower panels, respectively.

We next performed CIU experiments simultaneously on both F(ab’)_2_ and Fc subunits of adalimumab, panitumumab, and natalizumab after IdeS digestion (Figure 4). Overall, the differentiation of the mAb isotypes upon collisional activation of the F(ab’)_2_ domains is clearly evidenced based not only on the number of unfolding transitions observed in the CIU fingerprints, but also due to the different collision energies associated with each transition (Figure 4). While only two unfolding transitions are observed in the CIU fingerprint of the IgG2 F(ab’)_2_ domain (17.9 V and 37.6 V, respectively) (Figure 4b), three transitions are observed in the case of the IgG1 F(ab’)_2_ domain (23.6 V, 67.3 V, and 123.5 V respectively) (Figure 4a), and five transitions in the IgG4 F(ab’)_2_ CIU fingerprint (18.5 V, 72.5 V, 113.0 V, 147.6 V, and 162.4 V) (Figure 4c). The most unfolded state of the IgG2 (Fab’)_2_ domain is populated at lower voltages compare to the IgG1, and IgG4 isotypes. However, this final state is kinetically stabilized from 40 to 200 V, whereas the different unfolded states of IgG1, and IgG4 isotypes are only kinetically stabilized on shorter voltage ranges. The gas-phase stability of the F(ab’)_2_ domain of the IgG2 isotype stems more likely from the higher number of disulfide bridges in the hinge region (four inter-chain S-S) that prevents the unfolding process of the domain upon ion heating.

Conversely, CIU fingerprints of the Fc domains of the three mAb isotypes (adalimumab, panitumumab, and natalizumab) exhibit very similar unfolding patterns with two unfolding transitions that lead to an increase of the collision cross section, and a final transition around 120 V where the global conformation of the Fc domain is compacted (Figure 5). This result is consistent with the high similarity (~ 95%) of the Fc sequence between the three isotypes (Table S1) leading to very similar gasphase stabilities and dynamics. However, the collision energy associated to the unfolding transitions observed on the IgG1 Fc fingerprint (41.5 V, 86.4 V, and 131.5 V) are slightly higher compared to those observed in the IgG2 (25.0 V, 67.5 V, and 122.5 V) and IgG4 (27.7 V, 67.5 V, 117.4 V) Fc domains suggesting a slightly higher gas-phase stabilization of the IgG1 Fc domain. This observation might be related to the influence of the non-covalent interactions that contribute to the stabilization and dimerization of the mAb Fc domain^26–30^. Indeed, the strongest CH3-CH3 interaction was found in the IgG1 structure (up to 10^6^-fold) in comparison to the other isotypes^26^, which is in good agreement with the gas-phase stability observed in the Fc CIU fingerprints. To corroborate this hypothesis, the stability of the constant regions was also investigated using differential scanning calorimetry (DSC) (Table S2)^22^. In this case, the melting temperatures corresponding to the denaturation of the CH2 and CH3 domains of the different mAbs isotypes evidenced a higher thermal stability for the IgG1 heavy chain constant domains, in agreement with results obtained by middle-level CIU experiments.

Altogether, our results depict that middle level CIU patterns of F(ab’)_2_ domains generated after IdeS digestion enable more easy and reliable classification of mAb isotypes compared to intact level CIU. As expected, therapeutic mAb isotype differentiation based on CIU fingerprints of Fc domains seem to be less adequate since only very minor differences regarding the collision voltage associated with the unfolding transitions are observed.

### Middle level IM-MS strategies to uniquely tackle the “hybridicity” of IgG2/4 eculizumab

Eculizumab is a humanized hybrid IgG2/4 mAb directed against the complement protein C52 and indicated to treat the rare hemolytic disease paroxysmal nocturnal hemoglobinuria^31^. The heavy-chain constant region of the parental antibody was repaved with components of both human IgG2 and IgG4 constant regions. The heavy chain of the hybrid mAb includes the CH1 and hinge regions of human IgG2 fused to the CH2 and CH3 regions of human IgG4. To avoid the generation of an antigenic site during the fusion, a restriction endonuclease cleavage site common to both IgG2 and IgG4 was used to join the two constant regions (31 amino acids flanking the fusion site are identical between IgG2 and IgG4)^32^. The unique combination of an IgG2/4 constant region makes this molecule fail to bind to Fc receptors (IgG2) and does not activate complement cascade (IgG4), which reduces the pro-inflammatory potential of the antibody^33^. Due to its inherent hybrid constitution, classical analytical techniques applied to eculizumab characterization provide a series of unclear/contradictory results. For example, non-reduced capillary electrophoresis-sodium dodecyl sulfate (nrCE‐SDS) eculizumab analysis presents a single peak which is rather in agreement with an IgG4 than with an IgG2 nrCE-SDS behavior for which doublet peaks are expected (Figure S4)^34^. Conversely, reversed-phase high performance liquid chromatography (rpHPLC) analysis of eculizumab clearly presents an IgG2-like behavior with three peaks reflecting IgG2 structural isoforms A, B and A/B (Figure S5)^35–36^. As classical analytical methods are not adapted to depict and dissect the complex structural scaffold of hybrid mAb formats, there is a need for analytical techniques able to tackle this challenging issue. We thus applied our CIU workflows for eculizumab characterization.

Regarding IM-MS analysis and CCS measurements and as expected from our results on reference IgGs, hybrid eculizumab cannot be differentiated from reference therapeutic mAb isotypes using native IM-MS at the intact level (Figure S2). At the middle-level, independently of the charge state, the ^TW^CCS_N2_ of the (Fab’)_2_ region of eculizumab is closer to those of the IgG2 reference (panitumumab) compared to those of the IgG1 or IgG4 references (Figure 3), which is a first hint towards eculizumab (Fab’)_2_ region behaving as an IgG2. As expected, no conclusions can be drawn from middle level native IM-MS ^TW^CCS_N2_ measurements regarding the Fc part of eculizumab owing to the high primary sequence similarity (~95%) with the three mAbs (Table S1 and Figure S3). Altogether IM-MS investigation provides very limited information towards the characterization of the “hybridicity” of eculizumab.

We thus moved to CIU experiments. Intact-level CIU fingerprint of eculizumab (hybrid IgG2/IgG4) was compared to those of panitumumab and natalizumab (references IgG2, and IgG4, respectively) previously described (Figure 2d). Overall, eculizumab CIU fingerprint shows four unfolding states and three transitions, revealing one additional unfolding transition compared to the reference IgG2 or IgG4 isotypes. In more details, the first transition occurs at 36.6 V, the second at ~145 V and the last one at 168 V. Automated isotype classification using the CIUSUIte2 module^37^ does not lead to clear isotype classification even if it mainly recognizes eculizumab as an IgG2 but with a high RMSD (Fig. S6c). Interestingly, a closer manual data interpretation allowed highlighting that the first CIU transition of eculizumab (37 V) is similar to the first transition of IgG2 or IgG4, while the two other eculizumab transitions correspond to the second transition of reference IgG4 (145 V for eculizumab versus 147 V) or IgG2 (168 V), respectively, which might suggest that eculizumab CIU fingerprint could result from a composite/hybrid of the two IgG4 and IgG2 CIU patterns. This first evidence can be strengthened based on the centroid IM drift times of each unfolding state. The IM drift time of the second state of eculizumab is similar to the drift time of the second state of natalizumab (blue values in Figure 2i) while the drift time of the third state of eculizumab is close to the second unfolding state drift time of panitumumab (red values in Figure 2i). These data suggest that eculizumab gas-phase unfolding behaviour is hybrid between reference IgG2 and IgG4 mAbs. However, even if intact level CIU allows concluding that eculizumab CIU fingerprint is clearly different from reference IgG2/IgG4 ones, it does not allow to draw any conclusion about the origin of this difference related to its inherent “hybridicity”.

We finally performed CIU experiments on the F(ab’)_2_ and Fc subunits of eculizumab obtained upon IdeS digestion (Figure 4). Overall, the CIU fingerprint of the F(ab’)_2_ subdomain of eculizumab exhibits a very similar CIU pattern (same number of unfolding transitions at very similar collision energies) with the reference IgG2 F(ab’)_2_ (Figure 4b and d), suggesting an IgG2-like gas-phase unfolding of eculizumab F(ab’)_2_. Automatic isotype classification algorithm included in the open source CIUSuite2 software^19^ (Figure S6 d, e, and f) assessed the F(ab’)_2_ subdomain of eculizumab F(ab’)_2_ as an IgG2-type CIU pattern with less than 7.4 % of RMSD, which is a typical RMSD value between CIU replicates of the same mAbs^21, 23–24^. These results corroborate middle-IM-MS CCS measurements, and clearly show that the CIU pattern of eculizumab (Fab’)_2_ domain can be unambiguously associated to an IgG2 isotype.

For the Fc subdomains, as expected, very similar CIU patterns were observed for all mAbs (Figure 5). Automatic isotype detection of the CIUSuite2 software^19^ revealed that eculizumab Fc unfolding pattern was slightly closer to the Fc subdomain of the IgG4 reference (natalizumab) rather than the IgG1 or IgG2 ones (Figure 5, and S7). Indeed, the resulting RMSD upon comparison of the eculizumab Fc CIU fingerprint with both IgG2, and IgG4 references were 8.8 %, and 4.6 %, respectively, suggesting that the unfolding behavior of the Fc part of eculizumab resembles to an IgG4-like isotype. These subtle but significant differences were confirmed upon careful manual data interpretation. Differences between CIU patterns stem on the very close collision energies associated with each individual unfolding transitions observed in the eculizumab and IgG4 Fc fingerprints (27.7 V, 67.5 V and 117.8 V for eculizumab compared to 27.7 V, 67.5 V and 117.4 V for the reference IgG4). Conversely, all voltages associated to IgG1 unfolding pattern were significantly higher when compared to eculizumab, while only one voltage associated to the third unfolding event allows distinguishing eculizumab (117.8 V) from IgG2 (122.5 V) (Figure 5, and S7).

Altogether, our results clearly demonstrate that middle-level CIU is a unique MS-based approach to probe the duality/“hybridicity” of engineered mAbs formats. In our case, among all tested analytical techniques (nrCE-SDS, rpHPLC-UV, native IM-MS and CIU-IM-MS), middle-level CIU experiments was the only one able, within one single analysis run, to provide structural evidences of eculizumab hybrid format and to assess the “isotypicity” of each of its domains. The energy associated with the unfolding transitions along with the number of unfolding events present on F(ab’)_2_ and Fc CIU fingerprints afforded an accurate and straightforward identification of eculizumab hybrid construction.

## CONCLUSIONS

In this work, we highlight that middle-level IM-MS analyses (after IdeS digestion of mAbs), and more precisely middle-level CIU experiments, afford better differentiation of mAb isotypes than similar analyses performed at the intact level. At the intact level (150 kDa), mAb isotypes usually present “co-drifting”/overlapping ATDs when using commercially available IM-MS instruments due to a lack of IM resolution, which prevents isotype classification through a simple CCS measurement. Conversely, at the middle-level, IgG2 can be clearly distinguished from IgG1/IgG4 by the CCS measurements of its F(ab’)_2_, which is a first slight improvement.

More impressive conclusions for mAb isotype classification were obtained from middle-level CIU experiments, especially from F(ab’)_2_ CIU pattern interpretation (100 kDa). As the vibrational energy redistribution is more efficient upon collision of the smaller F(ab’)_2_ and Fc domains with the background gas, the ions in the gas-phase can populate additional excited unfolding states, which provide clear-cut specific signature characteristics of mAb isotypes. Conversely to CIU fingerprints of therapeutic mAbs recorded at the intact level that only present subtle differences in the 100-200 V region, F(ab’)_2_ CIU fingerprints exhibit significantly different unfolding features both in terms of number and associated energies of unfolding transitions throughout the whole voltage range (from 0 to 200 V). As a consequence, the F(ab’)_2_ CIU fingerprints can be considered as the most diagnostic region to differentiate mAb isotypes since the number of unfolding transitions and their associated energies are clearly different for IgG1, IgG2 and IgG4 mAbs. The unfolding behavior of the F(ab’)_2_ domains is directly related to isotype specific inter-chain disulfide connectivities that drive specific structures, leading to diagnostic CIU features. Although CIU unfolding patterns of Fc domains (50 kDa) are overall very similar owing to the absence of covalent connectivities in Fc domains (no inter-chain S-S bridges that connect Fc noncovalent dimers) and the high primary sequence similarities of Fc regions (> 93% in our study), minor differences can also be detected between middle-level CIU fingerprints of Fc domains. Indeed, a careful and detailed data interpretation of transition energies related to non-covalent dimeric Fc domains also affords distinguishing: i) IgG1 from IgG2 or IgG4, with overall higher unfolding energies for all transition states for IgG1 and ii) IgG2 from IgG4 on the basis of one unique transition (the more energetic at 117.4 V for IgG4 versus 122.5 V for IgG2). Ranking of gas-phase stabilities and resistance to unfolding of Fc non-covalent dimers (IgG1>IgG2>IgG4) were directly correlated to strength of noncovalent CH3-CH3 interactions. Our results thus show that middle CIU fingerprints are not only sensitive to covalent connectivities (disulfide bridges) differences that drive mAb structure and rigidity (F(ab’)_2_ domains) but also to non-covalent interactions (Fc domains).

Benefits of middle IM-MS and CIU approaches are clearly illustrated for the characterization of hybrid mAb-formats like the IgG2/IgG4 eculizumab. While classical analytical techniques such as nr-CE-SDS or rpHPLC-MS led to controversial results and failed in identifying the hybridicity of eculizumab, intact-level CIU approach provided a first strong hint towards a composite IgG2/IgG4 CIU pattern. At the middle-level, IM-MS analysis first revealed similar ^TW^CCS_N2_ values for the F(ab’)_2_ domains of eculizumab and the IgG2 mAb reference (panitumumab), highlighting the possibilities of CCS measurements to guide isotype classification at the middle-level. The precise IgG2/IgG4 “hybridicity” of eculizumab was definitely, more clearly and accurately assessed by middle-level CIU. Analysis of the middle-level CIU fingerprints of eculizumab pointed out that the F(ab’)_2_ unfolding pattern corresponds to an IgG2-like mAb reference, corroborating the results obtained using middle IM-MS analysis, while the Fc domain behaves as an IgG4-like isotype. For eculizumab, middle-level CIU experiments allowed uniquely to face the challenge of hybrid mAb-format characterization, allowing within one single CIU experiment to identify specific structural isotype features but also to attribute isotype to its corresponding mAb subdomain.

Altogether, our results highlight the suitability of middle-level CIU experiments to differentiate and classify the isotype of therapeutic mAbs, including complex new generation hybrid formats. Middle-level CIU provides more insights than intact-level CIU, which enables to overcome the limitation of classical analytical techniques. Even if progress in terms automation are still required to bring CIU to its mature state, the benefits associated with CIU at the middle-levels represent a real breakthrough therapeutic protein analysis, paving the way for CIU implementation in R&D laboratories.

## Supporting information

includes all supplemental figures and tables

## ASSOCIATED CONTENT

### Supporting Information

Supporting information Table S1, Table S2, Figure S1, Figure S2, Figure S3, Figure S4, Figure S5, Figure S6, and Figure S7 are available free of charge.

## Author Contributions

¶ (T.B., O.H.A.) These authors equally contributed to the work.

## Notes

The authors declare no competing financial interest.

## ACKNOWLEDGMENT

This work was also supported by the CNRS, the University of Strasbourg and the French Proteomic Infrastructure (ProFI; ANR10-INBS-08-03). The authors would like to thank GIS IBiSA and Région Alsace for financial support in purchasing a Synapt G2 HDMS instrument. T.B. and O.A.-H. acknowledge the Institut de Recherches Servier and the IdeX program of the University of Strasbourg for funding of their PhD and postdoctoral fellowship, respectively. E.D. was funded by the French Ministry of Higher Education, Research and Innovation for her PhD.

---

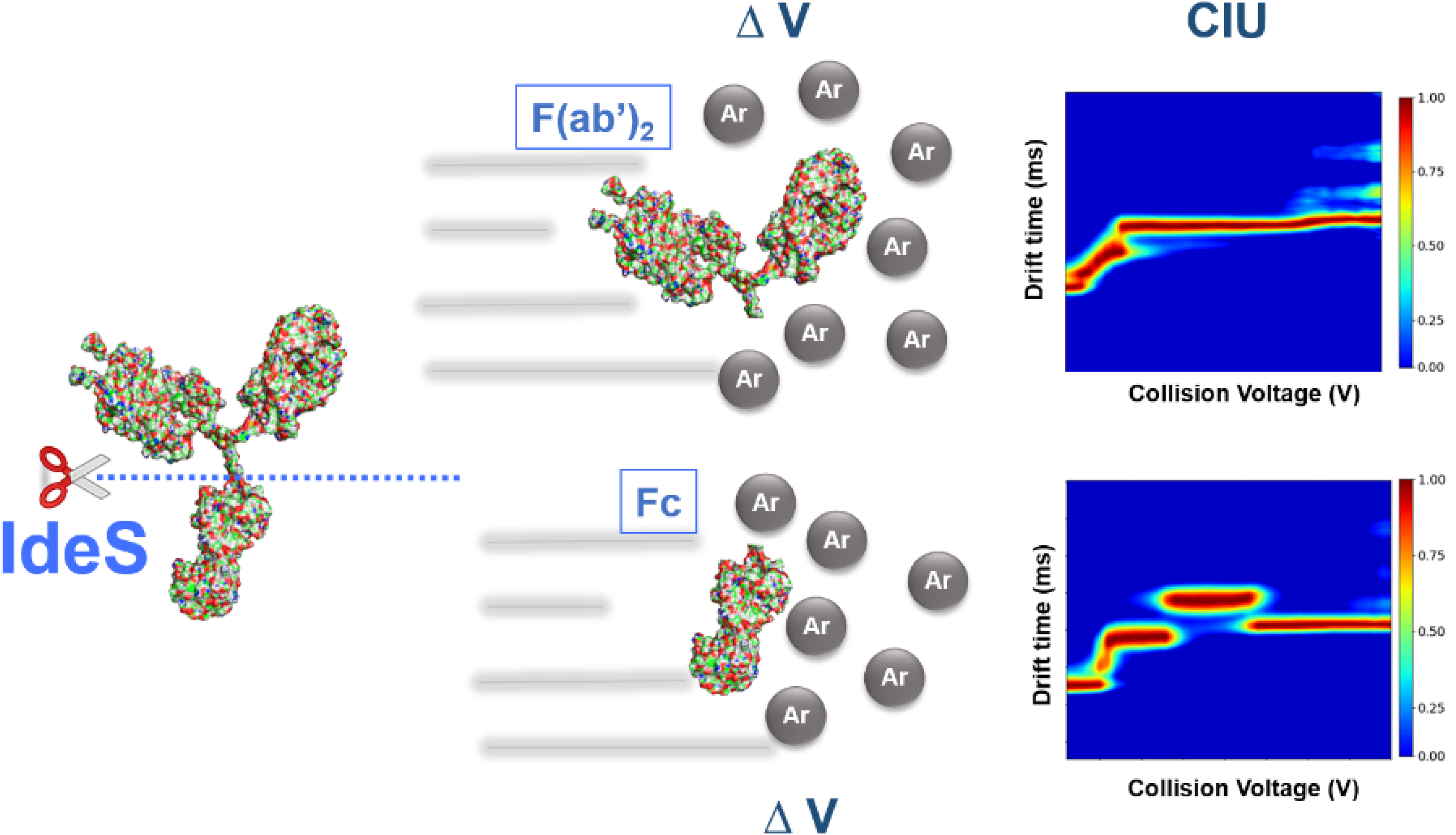

