## Supplementary material for "Middle-level IM-MS and CIU experiments for improved therapeutic immunoglobulin isotype fingerprinting": includes all supplemental figures and tables

### TABLE OF CONTENTS:

- Figure S1: Native mass spectra of adalimumab (a), panitumumab (b), natalizumab (c), and eculizumab (d).
- Figure S2: ATDs of 22+ charge state of adalimumab (IgG1), panitumumab (IgG2), natalizumab (IgG4) and eculizumab (IgG2/4) (a). Evolution of the $^{TW}CCS_{N2}$ of the intact mAbs as a function of the charge state (b). Table summarizing the measured $^{TW}CCS_{N2}$ of intact mAbs (c).
- Figure S3: Evolution of the Fc $^{TW}CCS_{N2}$ as a function of the charge state (a). Table summarizing the measured $^{TW}CCS_{N2}$ of the Fc domains.
- Table S1: Sequence homology of the F(ab')$_2$ (a) and Fc (b) domains of adalimumab (IgG1), panitumumab (IgG2), eculizumab (IgG2/4), and natalizumab (IgG4) mAbs.
- Table S2: Melting temperatures of CH2 and CH3 domains obtained from the DSC thermograms of different mAbs isotypes.
- Figure S4: Electropherograms of panitumumab (IgG2) (a), eculizumab (IgG2/4) (b), and natalizumab (IgG4) (c). Superposition of non-reduced capillary electrophoresis-sodium dodecyl sulfate (nrCE-SDS) runs with UV detection off.
- Figure S5: rpHPLC chromatogram (XIC) of adalimumab, eculizumab, and natalizumab.
- Figure S6: Differential CIU fingerprint between eculizumab and both panitumumab (IgG2) and natalizumab (IgG4) references of the 22+ charge state at the intact level (a, b), 21+ charge state of the F(ab')$_2$ domain (d, e), and 12+ charge state of the Fc regions (g, h). Isotype classification histograms provided by CIUSuite2 software are depicted in right panels (c, f, i).

- Figure S7: CIU gaussian fit of the 12+ charge state of Fc domains of panitumumab (a), eculizumab (b), and natalizumab (c). Differential CIU fingerprint between Fc domain of eculizumab and both IgG2 (d), and IgG4 (e) references. Isotype classification histogram of Fc domain of eculizumab based on the CIU data of the 12+ charge state (f).

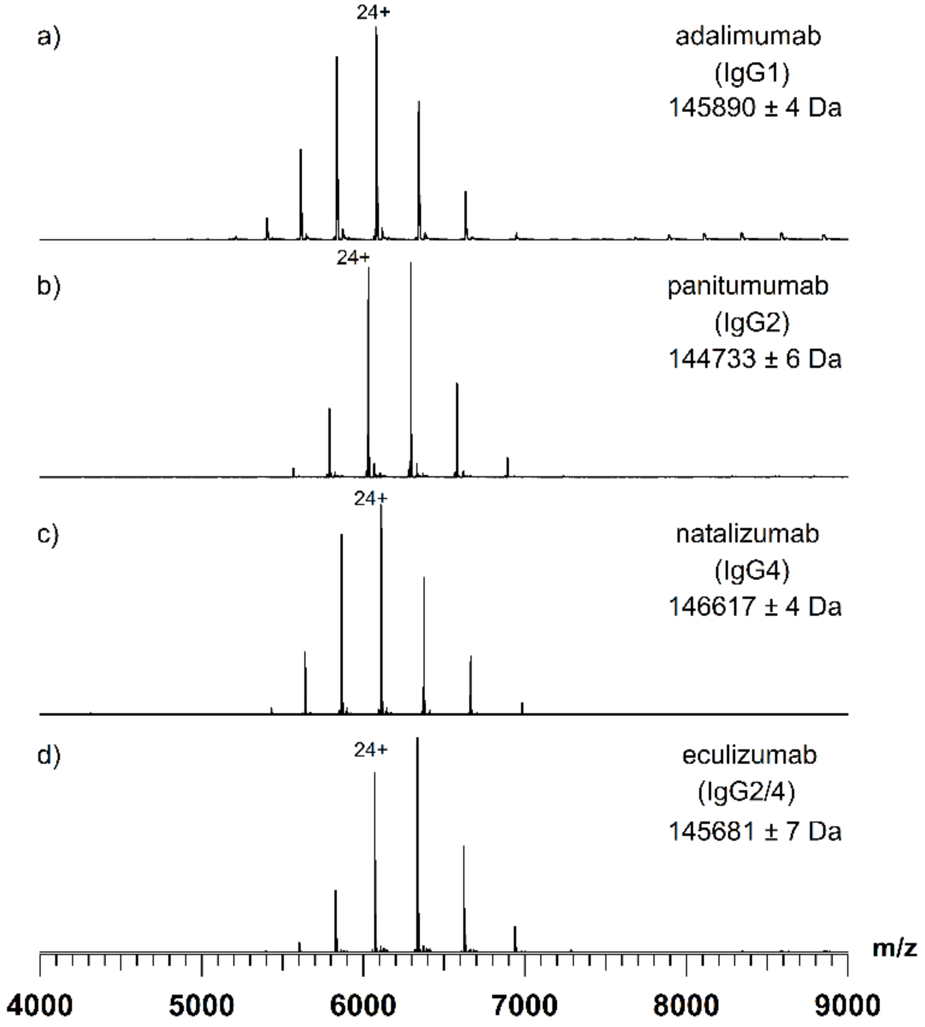


**Figure S1: Native mass spectra of adalimumab (a), panitumumab (b), natalizumab (c), and eculizumab (d).**

a)

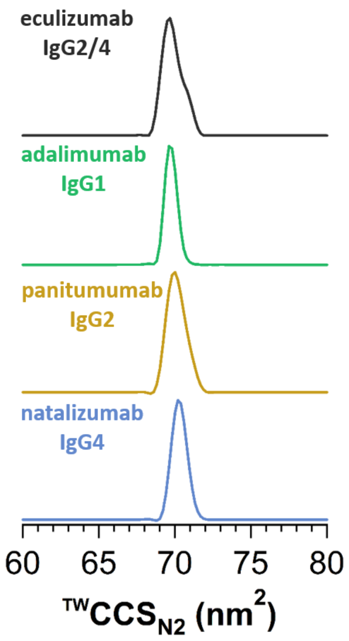


b)

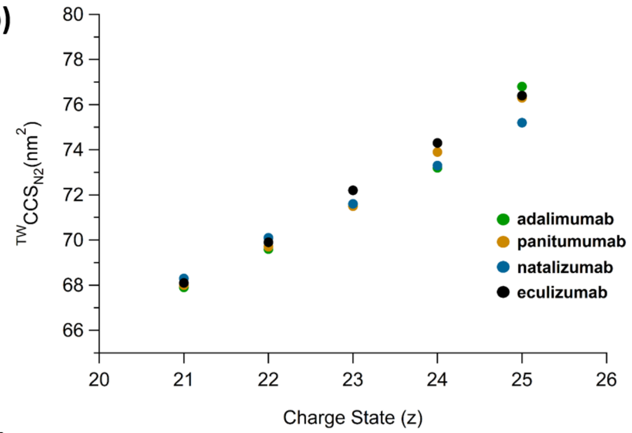


c)

| adalimumab | panitumumab | natalizumab | eculizumab |
|---|---|---|---|
| $^{TW}CCS_{N2}$(nm$^2$) | $^{TW}CCS_{N2}$(nm$^2$) | $^{TW}CCS_{N2}$(nm$^2$) | $^{TW}CCS_{N2}$(nm$^2$) |
| 67.9 ± 0.1 | 68.0 ± 0.1 | 68.3 ± 0.1 | 68.1 ± 0.1 |
| 69.6 ± 0.1 | 69.7 ± 0.1 | 70.1 ± 0.1 | 69.9 ± 0.1 |
| 71.5 ± 0.1 | 71.5 ± 0.1 | 71.6 ± 0.1 | 72.2 ± 0.1 |
| 73.2 ± 0.1 | 73.9 ± 0.1 | 73.3 ± 0.1 | 74.3 ± 0.1 |
| 76.8 ± 0.1 | 76.3 ± 0.1 | 75.2 ± 0.1 | 76.4 ± 0.1 |

**Figure S2: ATDs of 22+ charge state of adalimumab (IgG1), panitumumab (IgG2), natalizumab (IgG4) and eculizumab (IgG2/4) (a). Evolution of the $^{TW}CCS_{N2}$ of the intact mAbs as a function of the charge state (b). Table summarizing the measured $^{TW}CCS_{N2}$ of intact mAbs (c)**

**Table S1: Sequence homology of the F(ab')$_2$ (a) and Fc (b) domains of adalimumab (IgG1), panitumumab (IgG2), eculizumab (IgG2/4), and natalizumab (IgG4) mAbs**

| F(ab')$_2$ | adalimumab | panitumumab | eculizumab | natalizumab |
|---|---|---|---|---|
| **adalimumab** | | 77.1 % | 77.9 % | 76.8 % |
| **panitumumab** | 77.1 % | | 80.2 % | 75.2 % |
| **eculizumab** | 77.9 % | 80.2 % | | 81.5 % |
| **natalizumab** | 76.8 % | 75.2 % | 81.5 % | |

| Fc | adalimumab | panitumumab | eculizumab | natalizumab |
|---|---|---|---|---|
| **adalimumab** | | 95.7 % | 93.8 % | 94.3 % |
| **panitumumab** | 95.7 % | | 94.3 % | 94.8 % |
| **eculizumab** | 93.8 % | 94.3 % | | 99.5 % |
| **natalizumab** | 94.3 % | 94.8 % | 99.5 % | |

a)

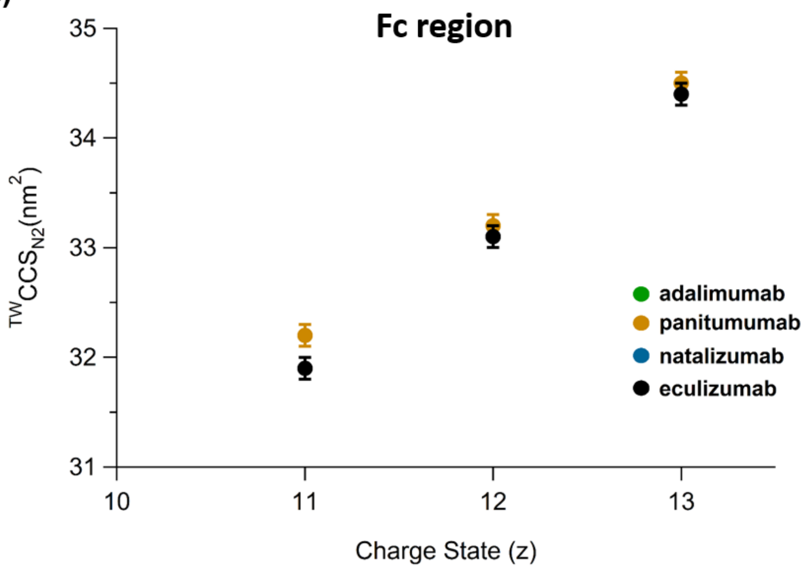


b)

| (z) | adalimumab $^{TW}CCS_{N2}(nm^2)$ | panitumumab $^{TW}CCS_{N2}(nm^2)$ | natalizumab $^{TW}CCS_{N2}(nm^2)$ | eculizumab $^{TW}CCS_{N2}(nm^2)$ |
|---|---|---|---|---|
| 11 | - | 32.2 ± 0.1 | 31.9 ± 0.1 | 31.9 ± 0.1 |
| 12 | 33.1 ± 0.1 | 33.2 ± 0.1 | 33.1 ± 0.1 | 33.1 ± 0.1 |
| 13 | 34.4 ± 0.1 | 34.5 ± 0.1 | 34.4 ± 0.1 | 34.4 ± 0.1 |
| * | 32.3 | 32.3 | 32.3 | 32.3 |

**Figure S3: Evolution of the Fc $^{TW}CCS_{N2}$ as a function of the charge state (a). Table summarizing the measured $^{TW}CCS_{N2}$ of the Fc domains (b).**

**Table S2: Melting temperatures of CH2 and CH3 domains obtained from the DSC thermograms of different mAbs isotypes**

| | Melting Temperatures (°C) | |
|---|---|---|
| | CH2 | CH3 |
| Adalimumab (IgG1) | 76.7 ± 0.1 | 83.2 ± 0.1 |
| Eculizumab (IgG2/4) | 68.7 ± 0.1 | 80.9 ± 0.1 |
| Panitumumab (IgG2) | 68.5 ± 0.1 | - |
| Natalizumab (IgG4) | 65.5 ± 0.1 | 80.8 ± 0.1 |

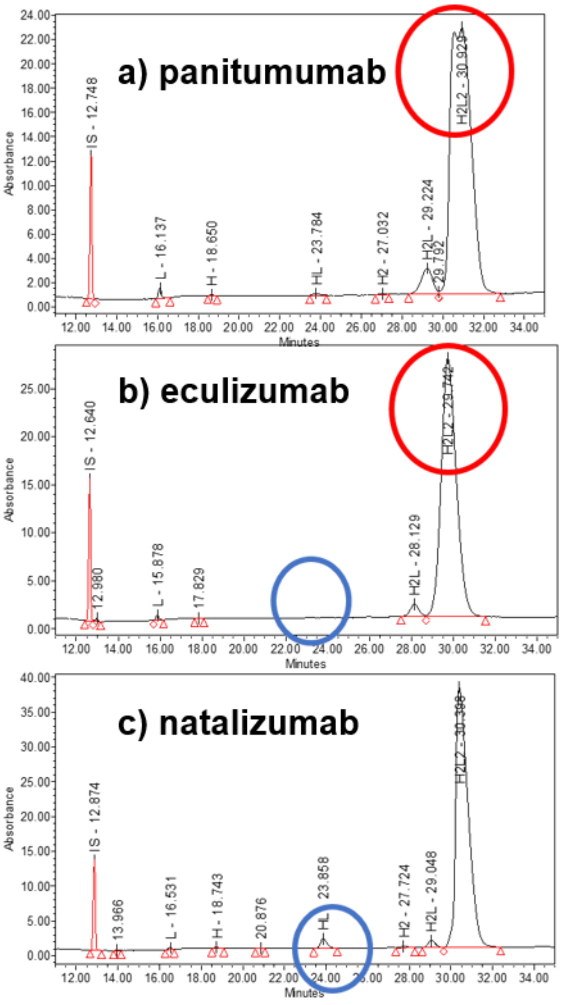


**Figure S4: Electropherograms of panitumumab (IgG2) (a), eculizumab (IgG2/4) (b), and natalizumab (IgG4) (c). Superposition of non-reduced capillary electrophoresis-sodium dodecyl sulfate (nrCE-SDS) runs with UV detection off.**

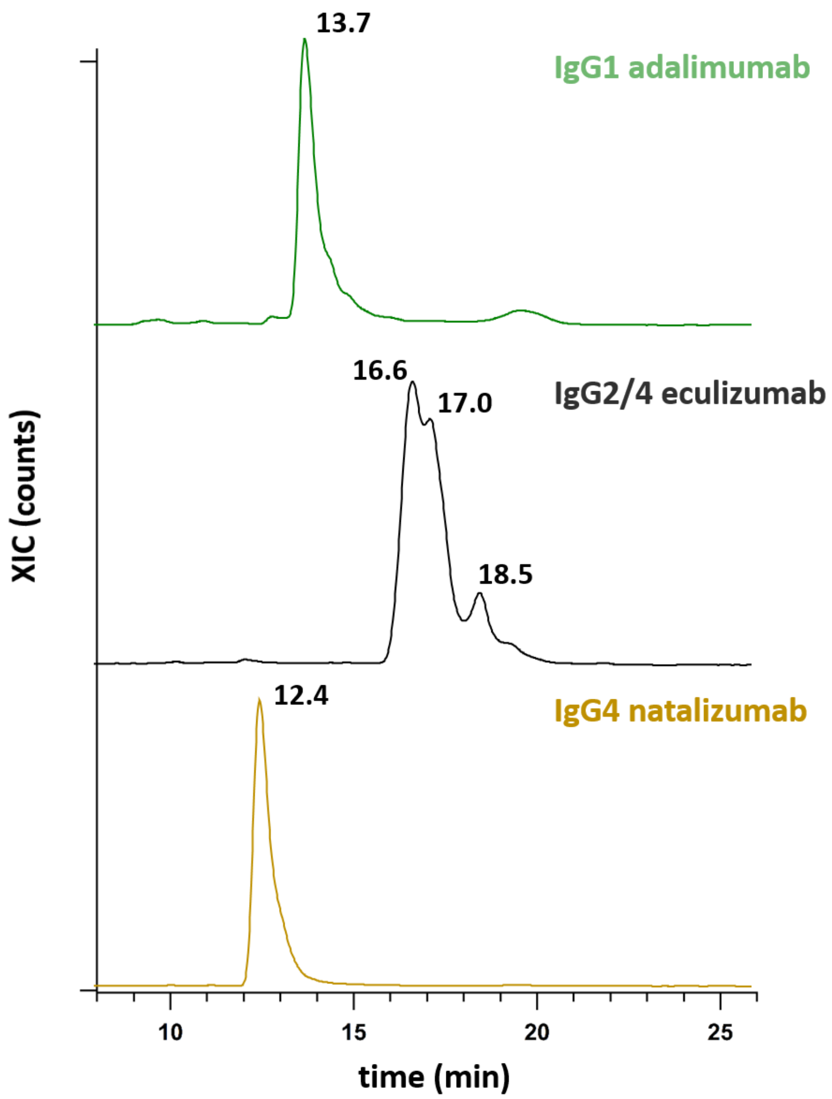


**Figure S5: rpHPLC chromatogram (XIC) of adalimumab, eculizumab, and natalizumab.**

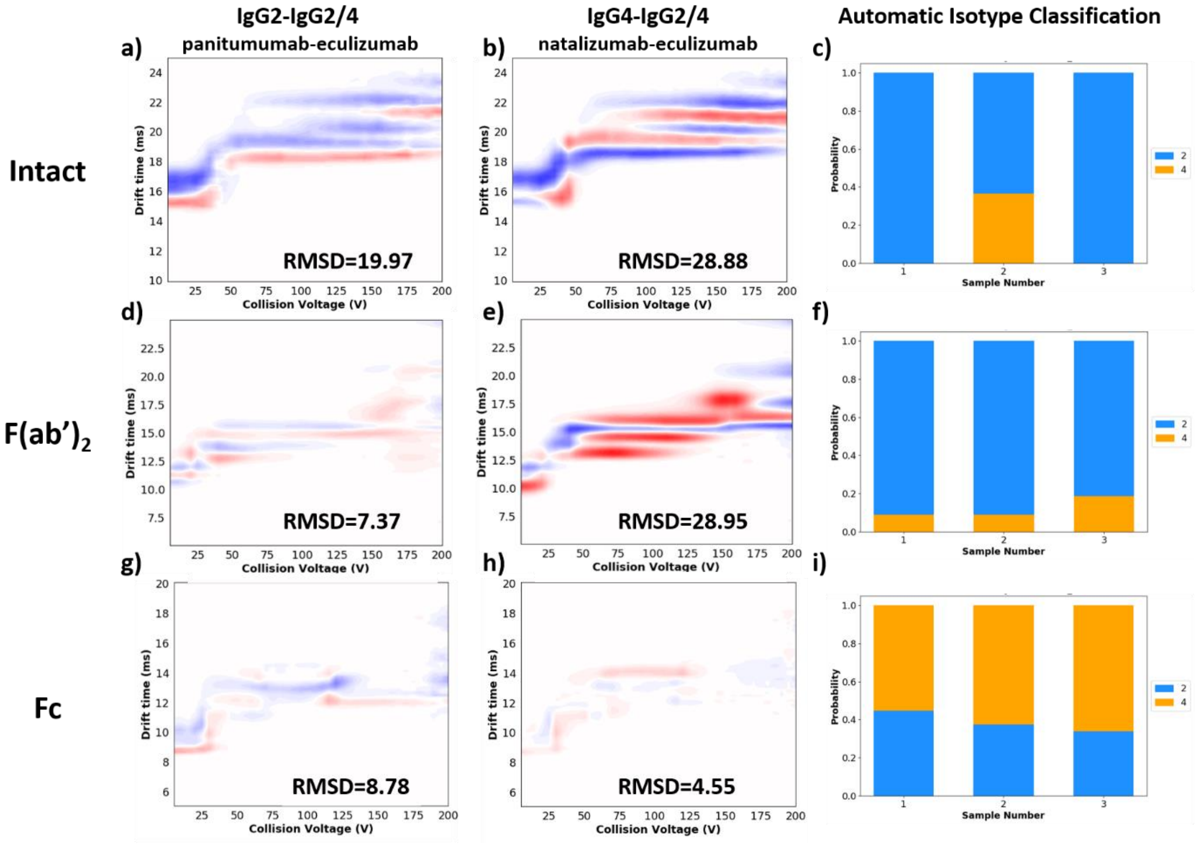


**Figure S6: Differential CIU fingerprint between eculizumab and both panitumumab (IgG2) and natalizumab (IgG4) references of the 22+ charge state at the intact level (a, b), 21+ charge state of the F(ab')$_2$ domain (d, e), and 12+ charge state of the Fc regions (g, h). Isotype classification histograms provided by CIUSuite2 software are depicted in right panels (c, f, i).**

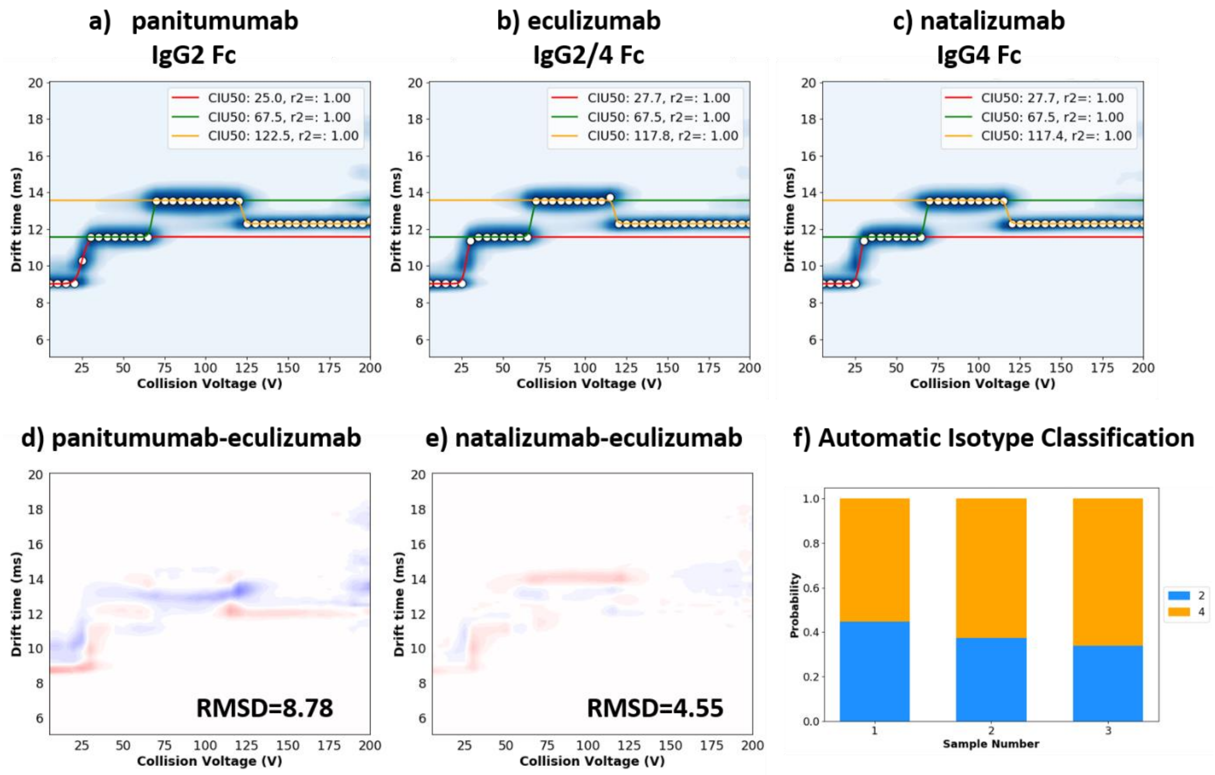


**Figure S7: CIU gaussian fit of the 12+ charge state of Fc domains of panitumumab (a), eculizumab (b), and natalizumab (c). Differential CIU fingerprint between Fc domain of eculizumab and both IgG2 (d), and IgG4 (e) references. Isotype classification histogram of Fc domain of eculizumab based on the CIU data of the 12+ charge state (f).**
